# Sphingosine-1-phosphate (S1P) signaling as a novel therapeutic target for alcohol abuse

**DOI:** 10.1101/2025.11.15.688635

**Authors:** Irene Lorrai, Riccardo Maccioni, Chenhao Wu, Chase Shankula, Jorge Marquez-Gaytan, Itzamar Torres, Roberta Puliga, Vez Repunte-Canonigo, Pietro Paolo Sanna

## Abstract

Sphingosine-1-phosphate (S1P) is a lipid mediator signaling through broadly expressed G protein-coupled receptors. We found that S1P is regulated by alcohol and that S1P receptor agonists reduce alcohol drinking in rodent models. Specifically, we observed that two S1P receptor agonists FDA-approved for multiple sclerosis, fingolimod and ozanimod, and the more brain penetrant S1P_1_ receptor agonist CYM5442, reduced binge alcohol drinking in the drinking in the dark (DID) paradigm in mice. CYM5442 also reduced drinking in dependent mice in the chronic intermittent ethanol vapor paradigm of dependence-induced increased drinking paired with 2 bottle-choice (CIE-2BC) as well as in non-dependent mice. CYM5442 reduced operant oral alcohol self-administration in both non-dependent and dependent rats made dependent by vapor exposure, and reduced motivation for alcohol in dependent rats tested in a progressive ratio schedule of reinforcement. CYM5442 significantly prevented cue-induced reinstatement in alcohol-dependent rats, a model of relapse to alcohol seeking. CYM5442 also reduced intake of non-drug reinforcers, including sucrose, food, water and, to a lesser extent, saccharine. Notably, CYM5442 was less aversive than naltrexone, an FDA-approved medication for the treatment of alcohol use disorder that shares a similar broad reducing action on alcohol intake and consummatory behavior. CYM5442 had no effect on loss of righting reflex, alcohol metabolism, motor coordination or spontaneous locomotor activity in rodents. Lastly, gene expression analysis by RNA-Seq revealed that S1P regulates a complex set of genes in the transition to alcohol dependence. Overall, our results establish S1P signaling as a novel therapeutic target for alcohol use disorder.

## Introduction

Alcohol use disorder (AUD) is a chronic, relapsing disorder characterized by the inability to stop or control alcohol use despite experiencing negative social, occupational, and health-related consequences. To date, only 3 medications have been approved by the Food and Drug Administration (FDA) for the treatment of AUD (disulfiram, oral and extended-release injectable naltrexone, and acamprosate) (1–3). Nalmefene has been approved in Europe and baclofen in France (1–3). Other medications such as topiramate, varenicline, ondansetron, gabapentin, aripiprazole, and prazosin/doxazosin have shown efficacy in some clinical trials (1–3). Due to AUD heterogeneity, both approved and investigational medications have limited efficacy, which reduces the confidence of clinicians in prescribing medications for AUD (3). Thus, there is pressing need to better understand the complexity of AUD and develop predicting models to guide drug selection to facilitate the utilization of existing AUD medications in clinical practice and to expand the therapeutic toolbox to better tailor AUD treatment to specific groups of patients and to identify broadly effective medications (3). The latter will require the identification and validation of new and more effective druggable targets (3).

Here we identified sphingosine-1-phosphate (S1P) signaling as a therapeutic target for AUD. S1P is a lipid mediator that affects multiple brain processes including neuroinflammation (4). S1P is derived from the phosphorylation of sphingosine by two sphingosine kinase (SphK) isoenzymes, SphK1 and SphK2, broadly expressed in the brain (5, 6). S1P can either act as a second messenger within the cells or can be released and signal through five G protein-coupled (S1P_1-5_) receptors (7) widely expressed in the body as well as in lymphocytes, neurons, astrocytes, oligodendrocytes, and microglia (8–11). S1P and S1P_1-5_ receptors are involved in physiological and pathological states including synaptic transmission, autophagy, and neuroinflammation, among others (7, 12, 13). The S1P receptors family has been identified as an important target for the treatment of chronic inflammatory states such as multiple sclerosis, ulcerative colitis, Crohn’s disease, and other conditions (14). Fingolimod, ozanimod and other S1P agonists are FDA-approved for relapsing-remitting multiple sclerosis and ozanimod also for ulcerative colitis.

Here, we investigated the role of S1P signaling in alcohol drinking and seeking. We observed that alcohol intake modulates S1P levels in the brain and that S1P agonists, including the FDA-approved medications fingolimod and ozanimod and the experimental compound CYM5442, reduce alcohol intake. These results establish S1P signaling as a therapeutic target for AUD.

## Material and Methods

### Animals

Male and female C57BL/6J mice (6 weeks old, The Jackson Laboratories, USA) were either single or group housed according to the experimental design. Male Wistar rats (4 weeks old, Charles River, USA) were housed in pairs. All animals were kept in standard plastic cages under controlled temperature (21 ±1°C) and humidity (50±5%). Food and water were available ad libitum, except when specified otherwise. Behavioral experiments were conducted during the dark phase of the light/dark cycle. All procedures adhered to the National Institutes of Health guidelines for the “Care and Use of Laboratory Animals” and were approved by the Institutional Animal Care and Use Committee of The Scripps Research Institute.

### Drugs

The S1P receptor agonists fingolimod, ozanimod, and CYM5442 (Tocris, MN, USA) were suspended in saline with 1% (w/v) Tween 80 and administered intraperitoneally (IP) 60 min prior to the behavioral experiments. The administration volumes were 10 ml/kg for mice and 2 ml/kg for rats.

### Experimental procedures

Details on the experimental procedures employed in the present study, including behavioral assays, targeted LC-MS/MS quantification of sphingosine-1-phosphate, total RNA Isolation, and RNA-Sequencing are provided in the Supplementary Material.

## Results

### Mice Experiments

#### Alcohol decreases S1P levels in the mouse prefrontal cortex

Targeted LC-MS/MS analysis was conducted to investigate the effects of alcohol on S1P levels. Male alcohol-naive C57BL/6J mice were administered with either saline or an intoxicating dose of alcohol [3.5 g/kg, 15% (w/v)]. Thirty minutes later the prefrontal cortex was collected and stored at −80°C until metabolomics analysis was performed. Statistical analysis revealed a significant decrease of S1P levels in the mouse group treated with alcohol (unpaired T-test, t_7_= 3.12; p<0.05) (Fig 1).

**Fig. 1.**
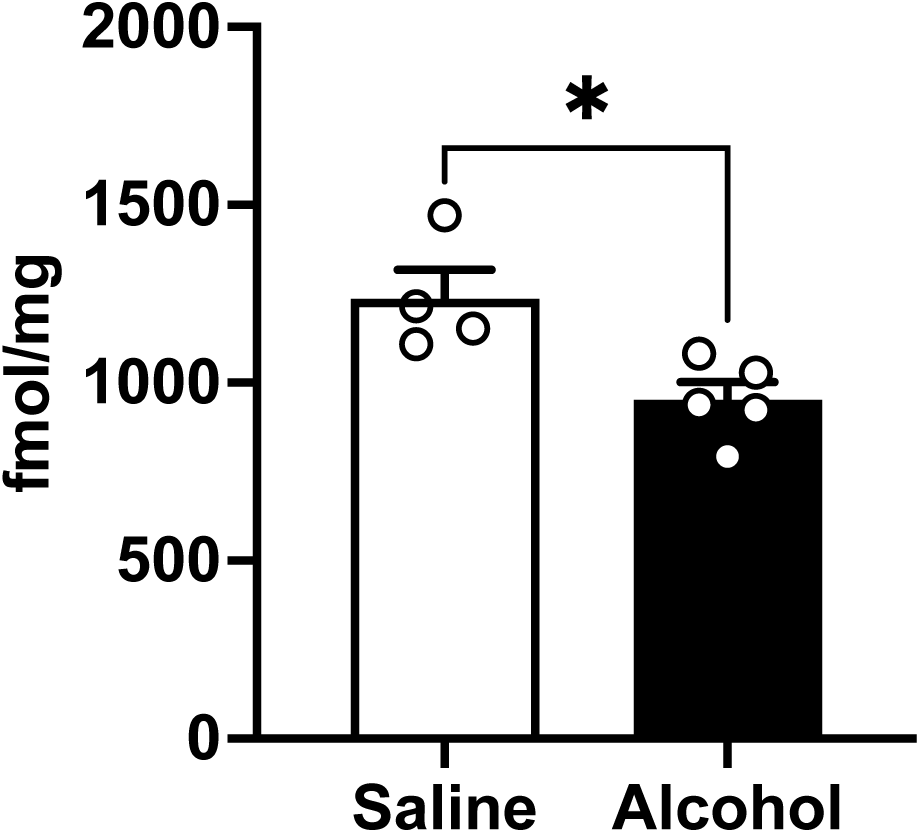
Alcohol reduces S1P levels in the Prefrontal Cortex of male C57BL/6J mice. Metabolomics of the prefrontal cortex (PFC) of C57BL/6J mice injected IP with alcohol (3.5 g/kg) or saline and euthanized 30 min after. Each bar represents the mean ± SEM of 4, 5 mice. (t_7_=3.12; p=0.017).

#### S1P receptor agonists reduces binge-like alcohol drinking

To evaluate whether pharmacological modulation of S1P signaling influences alcohol drinking, we examined the effect of S1P receptor agonists on binge-like alcohol drinking in C57BL/6J mice exposed to the drinking in the dark (DID) paradigm (15). We first assessed the effects of the two FDA-approved clinical compounds, fingolimod and ozanimod. Of note, fingolimod acts on almost all S1P receptor subtypes (S1P_1-5_) except S1P_2_, whereas ozanimod selectively targets S1P_1_ and S1P_5_. In contrast, CYM5442 is a highly selective agonist for the S1P_1_ receptor and has been shown to accumulate in the brain (16). Therefore, we focused subsequent studies on the pharmacological properties of CYM5442. These studies were also extended to include female mice.

Fingolimod. Acute administration of fingolimod in the dose-range between 1-4 mg/kg, effectively reduced alcohol consumption in the DID paradigm over both the first 2 h [1-way ANOVA, F(3, 50) = 11.41, p<0.0001] (Fig 2A) and the entire 4 h drinking session [1-way ANOVA, F(3, 50) = 16.74, p<0.0001; **\*\*\*\***p<0.0001 by Tukey’s post hoc test] in male C57BL/6J mice (Fig 2B).

**Fig. 2.**
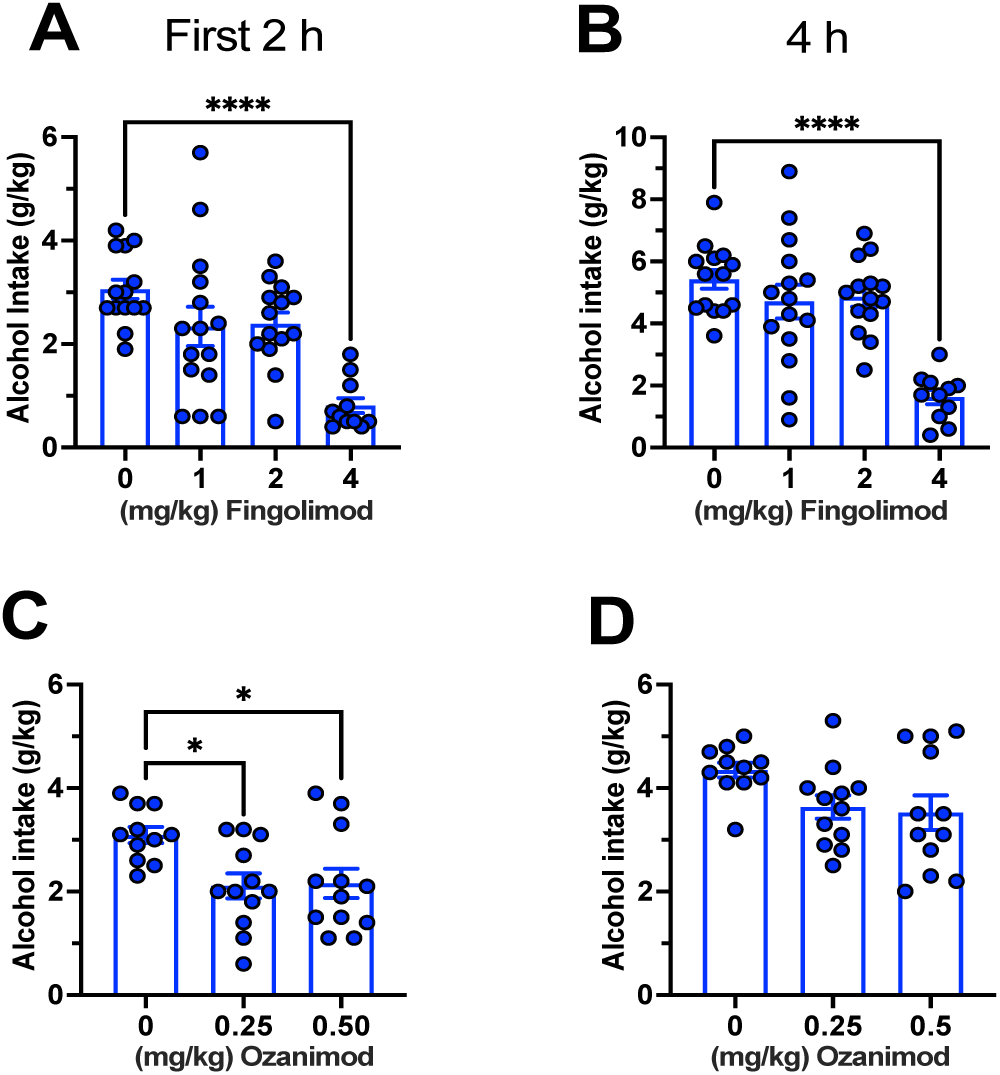
The S1P receptor agonists reduces alcohol intake in the drinking in the dark (DID) binge drinking paradigm in male C57BL/6J mice. *Fingolimod* (4 mg/kg) reduced alcohol consumption in the DID paradigm over the first 2 h **(A)** and the entire 4 h **(B)** drinking session in male C57BL/6J mice (**\*\*\*\***p<0.0001). Each bar represents the mean ± SEM of n=13-15 mice. *Ozanimod* (0.25 and 0.5 mg/kg) reduced alcohol consumption in the DID paradigm after the first 2 h **(C)** of drinking session in male C57BL/6J mice (p<0.05). The reducing effect of ozanimod was not evident at the end of the 4h drinking session **(D)**, which is consistent with its short half-life (172). Each bar represents the mean ± SEM of n=11-12 mice.

Ozanimod. Acute administration of ozanimod reduced alcohol consumption in the DID paradigm after 2 h in male C57BL/6J mice at both doses [1-way ANOVA, F (2, 32) = 5.33; p<0.05; **\***p<0.05 by Tukey’s post hoc test] (Fig 2C). However, the reducing effect of ozanimod was not evident at the end of the 4 h drinking session [1-way ANOVA, F (2, 32)=3.05, p>0.05] (Fig 2D), consistent with its short half-life (172).

CYM5442. We observed that acute administration of CYM5442 resulted in a statistically significant reduction of alcohol intake (1-way ANOVA: F(3,36) = 13.66; p<0.0001) in the mouse groups treated with 2.5 and 5 mg/kg in comparison to the mouse vehicle-treated group (**p<0.005; ****p<0.0001 by Tukey’s post hoc test) during the first 2 h of the drinking session (Fig 3A). Furthermore, the alcohol-reducing effect of CYM5442 persisted through the entire 4 h session at the highest administered dose [1-way ANOVA: F (3,36) = 17.46; p<0.0001; ****p<0.0001 by Tukey’s post hoc test] (Fig 3B). A similar pattern was observed in the female mice with an even stronger reduction of alcohol intake during the first 2 h session following administration of 2.5 and 5 mg/kg CYM5442 [1-way ANOVA: F (3,32) = 15.51; p<0.0001; ****p<0.0001 by Tukey’s post hoc test] (Fig 3C). At the end of the 4 h session, alcohol consumption remained significantly reduced at the highest dose [1-way ANOVA: F (3,32) = 6.30; p<0.0005; **p<0.005, by Tukey’s post hoc test] (Fig 3D).

**Fig. 3.**
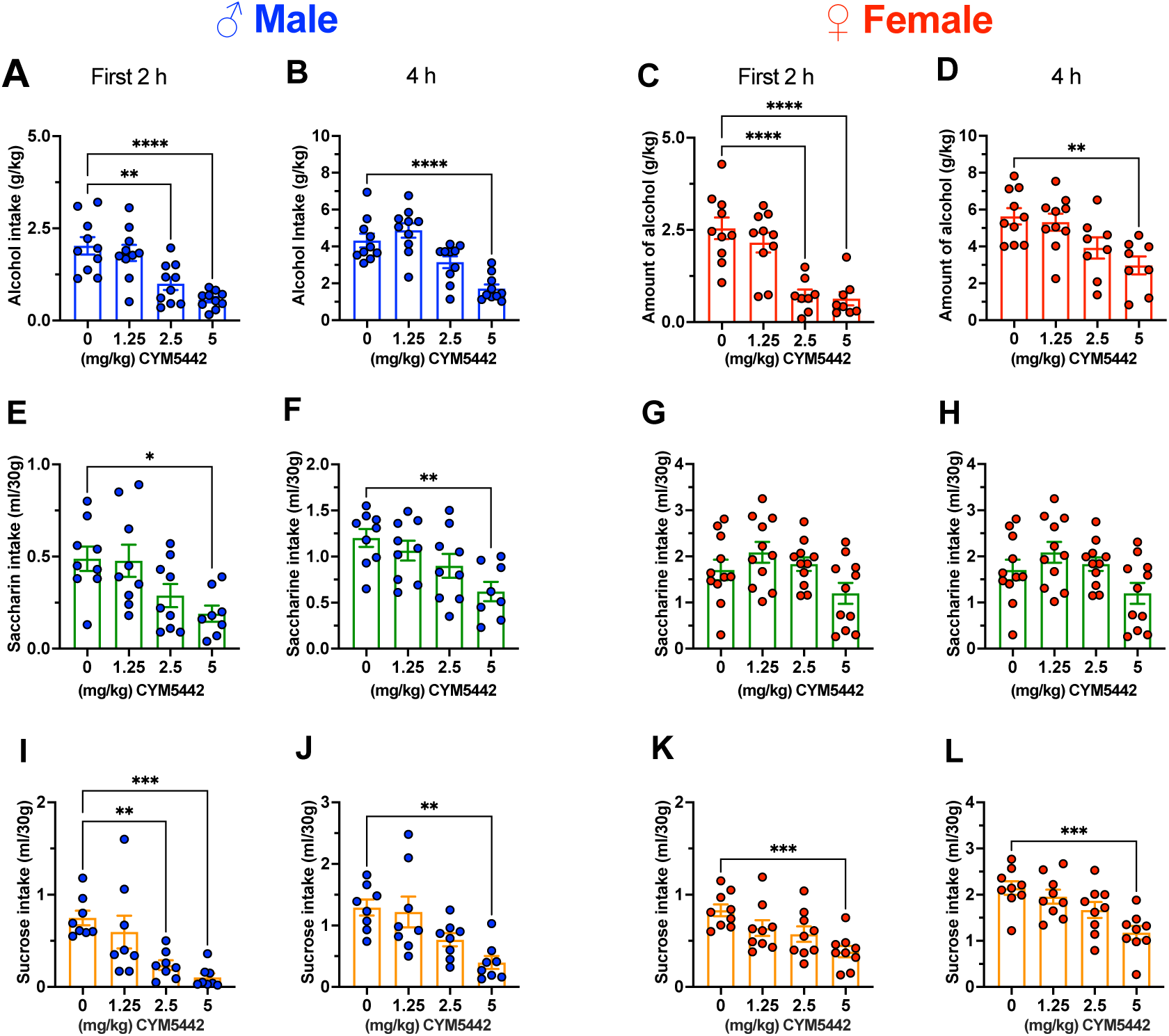
The selective S1P receptor 1 agonist CYM5442 reduces alcohol, saccharin, and sucrose intake in the DID binge drinking paradigm in male and female C57BL/6J mice. CYM5442 (2.5 and 5 mg/kg) reduced alcohol intake in the DID paradigm over the first 2 h **(A, C)** and the entire 4 h drinking session **(B, D)** in male and female C57BL/6J mice (**\*\***p<0.005; **\*\*\*\***p<0.0001). Male experiment: each bar represents the mean ± SEM of 10 mice. Female experiment: each bar represents the mean ± SEM of 8-10 mice. CYM5442 (5 mg/kg) reduced saccharin [0.002% (w/v)] intake in the DID paradigm over the first 2 h in male **(E)** but not female **(G)** mice. A similar pattern was observed at the end of the entire 4 h drinking session (*p<0.05, **p<0.005) **(F, H)**. Male experiment: each bar represents the mean ± SEM of 8-9 mice. Female experiment: each bar represents the mean ± SEM of 11 mice. CYM5442 (2.5 and 5 mg/kg) reduced sucrose [0.05% (w/v)] consumption in the DID paradigm over the first 2 h **(I, K)** and entire 4 h session **(J, L)** in both male and female mice. (**p<0.005, ***p<0.0005) **(F, H)**. Male experiment: each bar represents the mean ± SEM of 8 mice. Female experiment: each bar represents the mean ± SEM of 9 mice.

#### CYM5442 reduces non-drug reinforcers intake in the drinking in the dark paradigm in male and female C57BL/6J mice

To better delineate CYM5442 specificity, we tested it on two palatable non-alcohol liquid reinforcers with different caloric value, saccharine and sucrose, in male and female mice in a DID-like experimental design. We found that in male mice, saccharine intake (ml/30g) was slightly reduced following administration of CYM5442 during the first 2 h session (1-way ANOVA: F (3,31) = 4.51; p<0.05). Tukey’s post hoc analysis indicated that the 2.5 and 5 mg/kg doses produced statistically significant reductions (**p<0.005 and *p<0.05, respectively). (Fig 3E). By the end of the 4 h session, a significant decrease in saccharine intake remained evident at the 5 mg/kg dose [1-way ANOVA F (3,31) = 4.88; p<0.01; ****p<0.01 by Tukey’s post hoc test] (Fig 3F). In female mice, CYM5442, at the highest dose of 5 mg/kg, significantly reduced saccharine intake during the first 2 h session (1-way ANOVA F (3,40) = 5.83; p<0.005; *p<0.05 by Tukey’s post hoc test) (Fig 3G). At the end of the entire 4h a slight reduction in saccharin intake was still observed (1-way ANOVA F (3,40) = 3.23; p<0.05), however, post hoc analysis did not reveal any statistically significant differences between groups (Fig 3H).

A similar pattern was observed with sucrose intake. Specifically, in male mice, administration of CYM5442 reduced sucrose intake (ml/30g) during the first 2 h session (1-way ANOVA: F (3,28) = 8.62; p<0.0005), particularly at the 2.5 and 5 mg/kg doses, as indicated by Tukey’s post hoc test (**p<0.01 and ***p<0.001, respectively) (Fig 3I). At the end of the entire 4 h session, only the highest dose (5 mg/kg) produced a significant reduction in sucrose intake (1-way ANOVA: F (3,28) = 6.85; p<0.005; **p<0.005 by Tukey’s post hoc test) (Fig 3J). In female mice, only the highest dose of CYM5442 (5 mg/kg) significantly reduced mice sucrose intake during both the first 2 hours (1-way ANOVA F (3,32) = 6.16; p<0.005; ***p<0.001 by Tukey’s post hoc test) (Fig 3K) and the entire 4 h session (1-way ANOVA F (3,32) = 7.41; p<0.005; ***p<0.001 by Tukey’s post hoc test) (Fig 3L).

#### CYM5442 reduces alcohol drinking in 2-bottle choice (2BC) after chronic intermittent ethanol (CIE) in male and female C57BL/6J mice

The effects of CYM5442 were also evaluated in the chronic intermittent ethanol vapor paradigm of dependence-induced increased drinking paired with 2 bottle-choice (CIE-2BC) (17, 18) in male and female mice. In male mice, two-way RM ANOVA of alcohol intake (g/kg) during the limited 2BC session revealed a significant main effect of alcohol vapor exposure (F (1,17) = 7.74; p<0.05) and treatment (F (1,17) = 13.97; p<0.005) as well as significant interaction (F (1,17) = 8.30; p<0.05). As expected, post hoc analysis indicated a significant difference in alcohol intake between the two control groups, with the dependent mice consuming an average of 4.30 g/kg and the non-dependent mice consuming an average of 1.50 g/kg (**p<0.005 by Šidák post hoc test) (Fig 4A). Administration of CYM5442 (5 mg/kg), drastically reduced alcohol intake in both groups (0.42 g/kg vs 1.15 g/kg, respectively). However, statistical significance was reached only in the dependent group (***p< 0.0005, Šidák post hoc test) (Fig 4A). In female mice, two-way RM ANOVA of alcohol intake (g/kg) during the limited 2BC session revealed a significant main effect of alcohol vapor exposure (F (1,18) = 4.74; p<0.05) and treatment (F (1, 18) = 36.23; p<0.0001) with a trend toward a significant interaction (F (1,18) = 3.75; p=0.06) (Fig 4B). Post hoc analysis showed that control dependent mice consumed more alcohol than non-dependent mice (6.11 g/kg vs. 4.11 g/kg, respectively). Administration of CYM5442 significantly reduced alcohol consumption in both non-dependent (*p< 0.05) and dependent (****p<0.0001) groups compared to their vehicle-treated control groups (Šidák post hoc test) (Fig 4B).

**Fig. 4.**
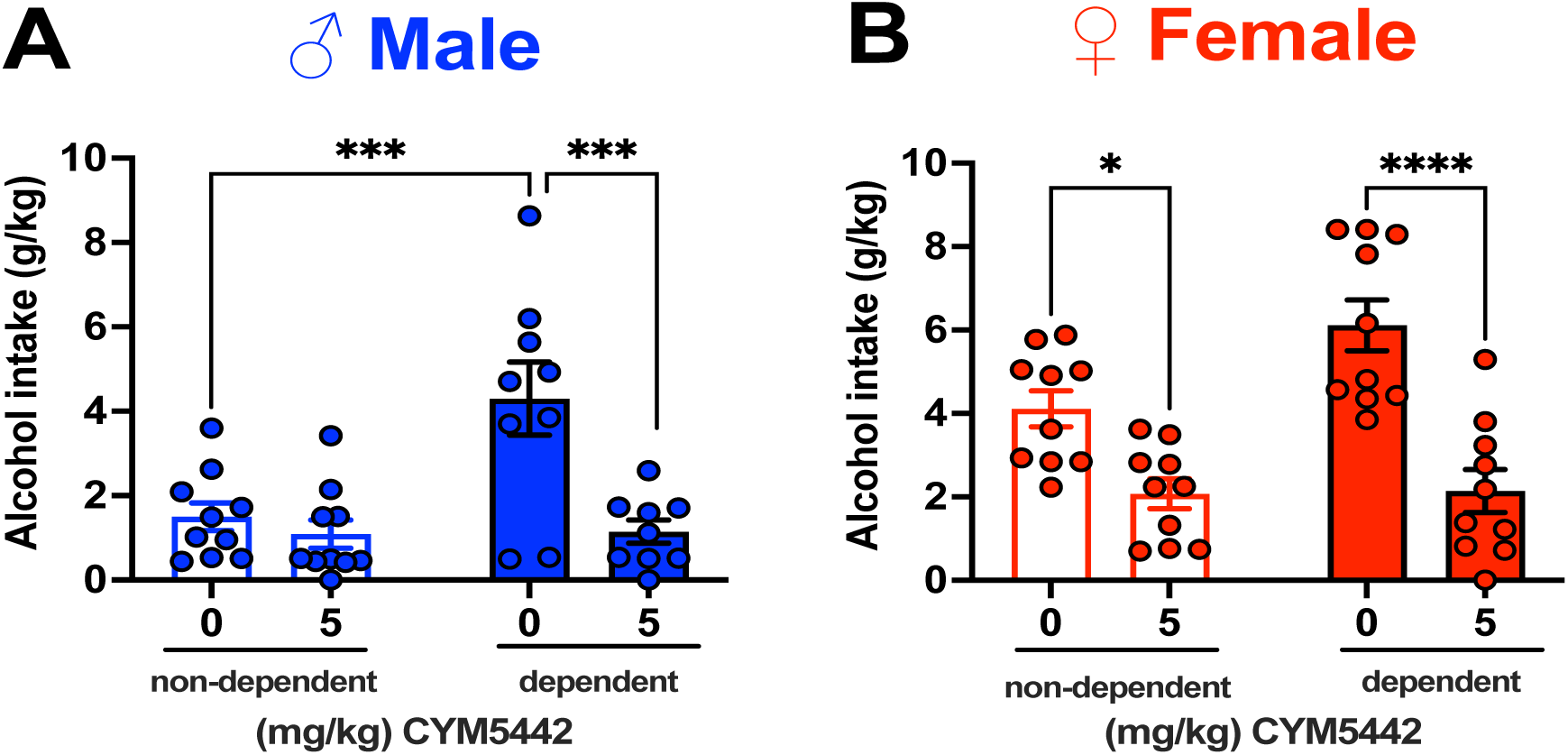
The selective S1P receptor 1 agonist CYM5442 reduces alcohol intake in non-dependent and dependent mice. **A)** Male dependent vehicle-treated mice consume more alcohol than non-dependent vehicle-treated mice as consequences of the chronic intermittent ethanol exposure with 2 bottle choice testing (CIE-2BC). Administration of CYM5442 effectively reduced alcohol intake in both groups; however statistical significance was reached only in the dependent group (***p <0.0005). Each bar represents the mean ± SEM of non-dependent (n=10) and dependent (n=8-9) mice. **B)** Similarly, female dependent vehicle-treated mice consume more alcohol than non-dependent vehicle-treated mice following CIE exposure, although no statistical significance was reached. CYM5442 administration significantly reduced alcohol intake in both groups, with a more pronounced effect observed in the dependent group (**\*\*\*\***p<0.0001), than non-dependent group (*p<0.05). Each bar represents the mean ± SEM of non-dependent (n=10) and dependent (n=10) mice.

#### CYM5442 reduces food, water intake, and energy metabolism in male C57BL/6J mice

CYM5442 effect was also tested in feeding, water intake and energy metabolism in male alcohol naive mice. Statistical analysis revealed that food and water intake were reduced by administration of CYM5442 over both the first 2 [food: 1-way ANOVA (F(3,26) = 7.87; p=0.0007), water (F(3,26) = 8.155; p=0.0005) (Fig 5A, C) and the entire 4 h session (food: 1-way ANOVA F(3, 26) = 3.014; p<0.05), water (F(3,26) = 3.147; p<0.05)), (*p<0.05; **p<0.01; ***p<0.0005; Tukey’s post hoc) (Fig 5B, D). In addition, CYM5442 significantly decreased mouse respiratory exchange ratio (RER) over both the first 2 h [1-way ANOVA F(3,26) = 4.22; p<0.05) (Fig 5E) and 4 h session (1-way ANOVA F(3,26) = 3.14; p<0.05), although only at the highest dose of 5 mg/kg (*p<0.05; **p<0.01, Tukey’s post hoc) (Fig 5F).

**Fig. 5.**
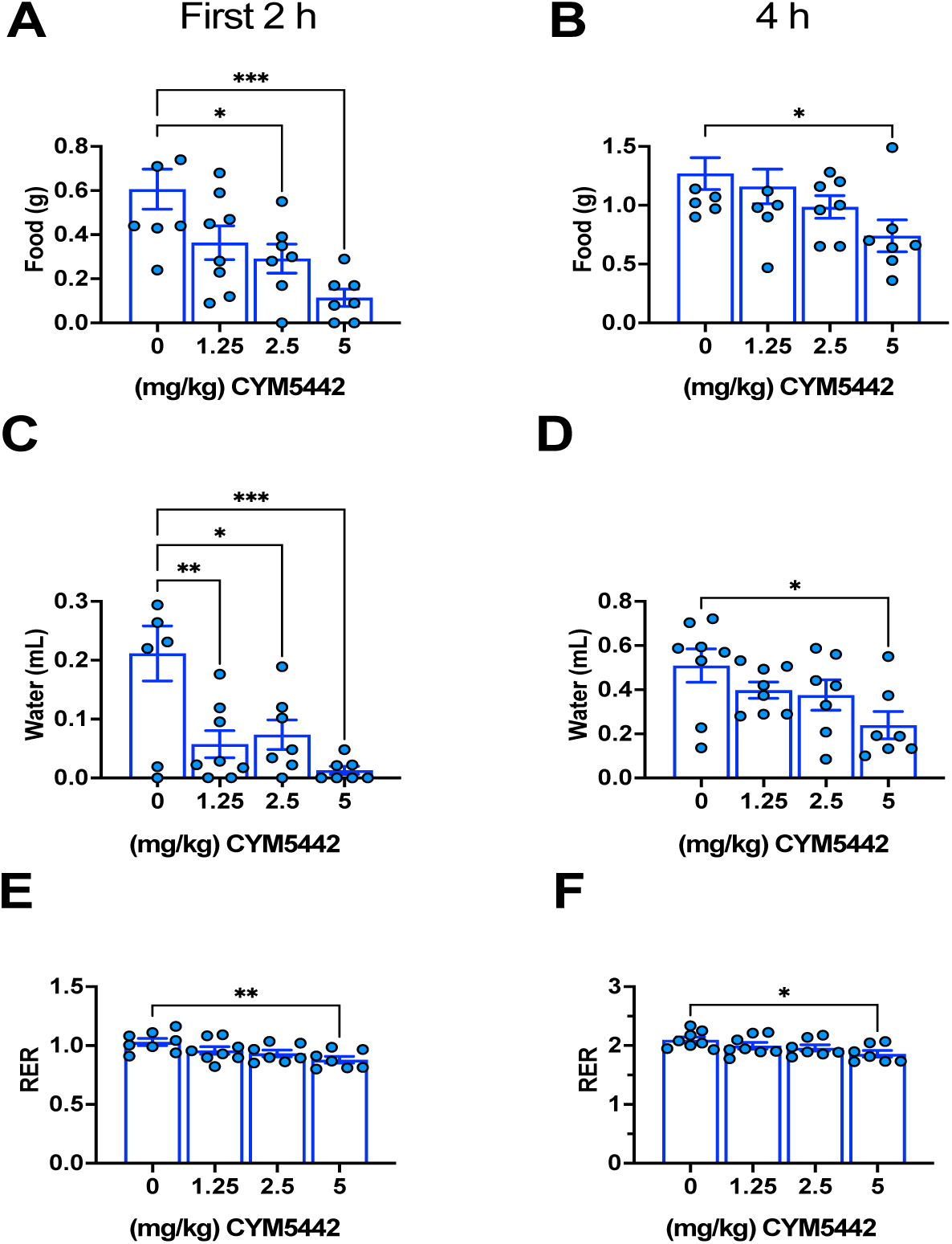
The selective S1P receptor 1 agonist CYM5442 reduces food, water, and respiratory exchange ratio in male mice. CYM5442 reduced water and food intake over both the first 2 h (Fig 5A, C) and the entire 4 h session (Fig 5B, D) (*p<0.05; **p<0.01; ***p<0.0005; Tukey’s post hoc). Similarly, CYM5442 reduced respiratory exchange ratio (RER) over both the first 2 h (Fig 5D) and the entire 4 h session, although this effect was significant only at the highest dose (5 mg/kg) (*p<0.05; **p<0.01, Tukey’s post hoc) (Fig 5E).

#### CYM5442 is less aversive than naltrexone in the conditioned place aversion paradigm

Alcohol-naive male mice were tested in the conditioned place aversion paradigm to evaluate potential aversive effects of CYM5442. A 2-way RM ANOVA revealed a significant main effect of time (F (1,26) = 16.03, p<0.001), but not treatment, although a trend toward significance was observed (F (2,26) = 2.96, p=0.07). In addition, a significant treatment x time interaction (F (2,26) = 6.71, p<0.005) was also found. Post hoc analysis showed a highly significant difference between pre and post conditioning in the naltrexone-treated group (***p=0.0001, by Šidák post hoc test) and a moderately significant difference was observed in the CYM5442-treated group (p=0.045, by Šidák post hoc test) (Fig 6). Overall, these data suggests that naltrexone produces a stronger aversive effect compared to CYM5442.

**Fig 6.**
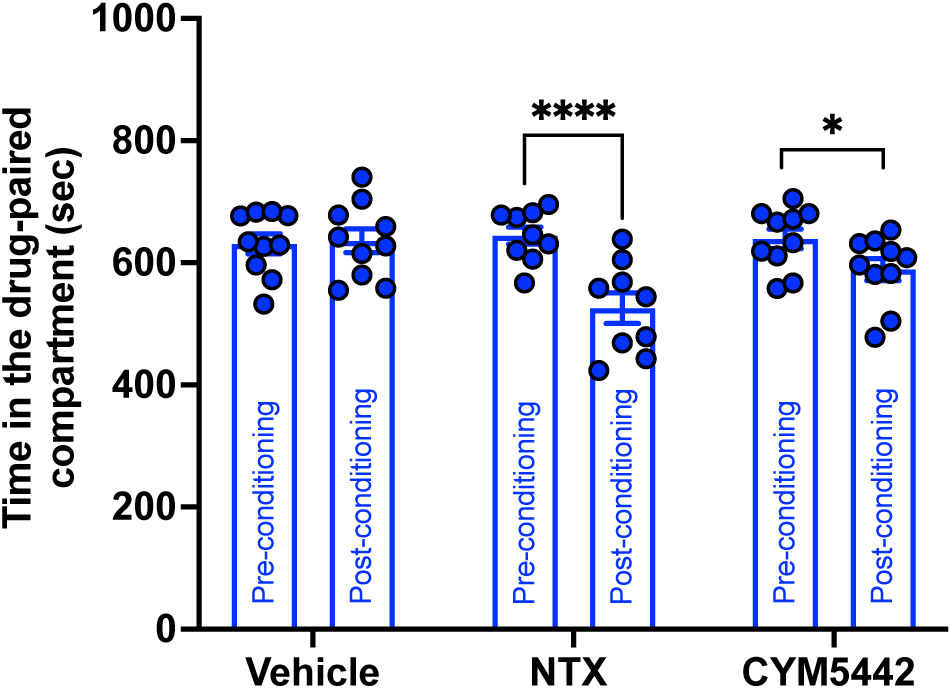
The selective S1P receptor 1 agonist CYM5442. Is less aversive than naltrexone (NTX) in male C57BL/6J mice as assessed by the conditioned place aversion test (*<0.05; **\*\*\***p<0.0001). Each bar represents the mean ± SEM of n=9-10 mice.

#### CYM5442 does not affect loss of righting reflex (LORR) or alcohol metabolism in male and female mice

To determine whether CYM5442 affects mouse sensitivity to the sedative and hypnotic effects of alcohol, male and female alcohol-naive mice were tested in the LORR paradigm. Statistical analysis indicated that duration of LORR was comparable between male mice treated with either 0 or 2.5 mg/kg CYM5442 and no significant difference was observed (unpaired two-tailed t-test, t_26_ = 1.38, p>0.05) (Fig 7A). Similarly, in females, treatment had no significant effect on LORR duration (unpaired two-tailed t-test, t_29_ = 1.05, p>0.05) (Fig 7D). Additionally, separated 2-way RM ANOVA of the BALs over time revealed a significant main effect of time in both sexes, but not significant effect of treatment or time x treatment interaction. For males: Time F (3,84) = 1401; p<0.0001; Treatment F (1,28) = 1.26; p>0.05; Interaction F (3,84) = 1.27; p>0.05) (Fig 7B). For females: 2-way RM ANOVA, Time F (3,90) = 2117; p<0.0001; Treatment F (1,30) = 0.002; p>0.05; Interaction F (3,90) = 2.17; p>0.05) (Fig 7E).

**Fig 7.**
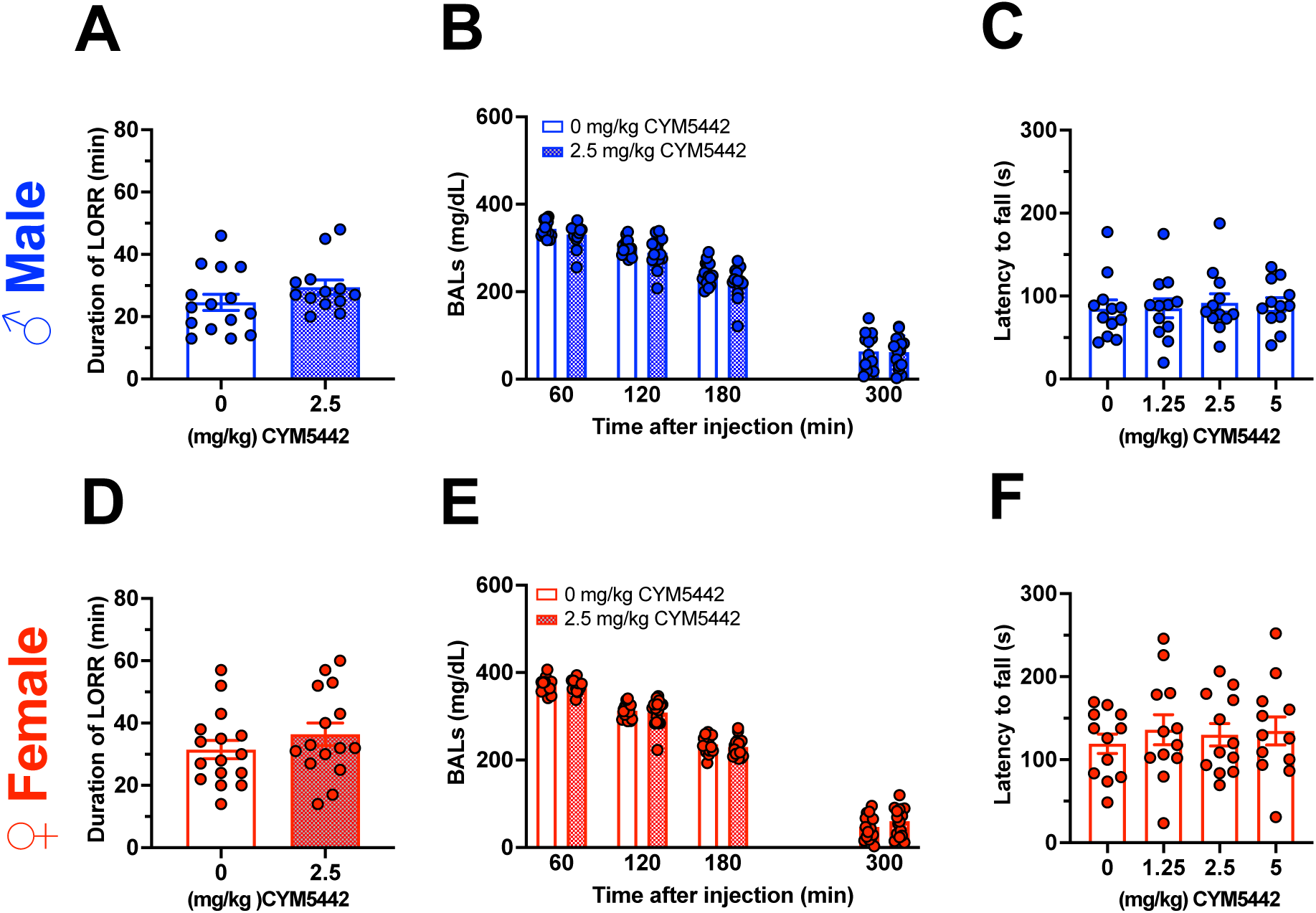
The selective S1P receptor 1 agonist CYM5442 does not alter mouse sensitivity to the alcohol hypnotic/sedative effects nor alter mouse motor coordination. CYM5442 (2.5 mg/kg) did not alter duration of LORR nor alcohol metabolism in male **(A, D)** or female **(B, E)** mice. Moreover, no dose of CYM5442 altered mouse performance on the rotarod apparatus (**C, F)**. Each bar represents the mean ± SEM of n=12 mice.

#### CYM5442 does not alter mouse motor coordination

Potential unspecific effects of CYM5442 were assessed using the rotarod apparatus in both male and female alcohol-naive C57BL/6J mice. Administration of CYM5442 had no effect on motor performance in either sex. Accordingly, 1-way ANOVA revealed no significant differences in latency to fall among treatment groups for males [F (3,44) = 0.11; p>0.05] (Fig 5C) or females (F (3,44) = 0.25; p>0.05) (Fig 7F).

### Rat experiments

#### CYM5442 reduces alcohol self-administration on a fixed ratio 1 (FR1) and a progressive ratio (PR) schedule of reinforcement in non-dependent and dependent Wistar rats

We then investigated whether the ability of CYM5442 to reduce binge-like alcohol drinking in mice, extended to a well-established operant paradigm of alcohol self-administration on FR1 in non-dependent and dependent male Wistar rats. Statistical analysis revealed that alcohol dependent rats exhibited significantly higher responding on the alcohol lever compared to non-dependent rats (63.6 vs 37.0 lever presses over 30-min session; main effect of group: F (1,103) = 20.66; p<0.0001). Two-way ANOVA revealed that CYM5442 administration significantly reduced alcohol self-administration in both non-dependent and dependent rat groups (main effect of treatment: F (3, 103) = 18.78; p<0.0001). No significant interaction between treatment and group was observed (F (3,103) = 1.467; p>0.05). Post-hoc analysis indicated that only the10 mg/kg dose of CYM5442 significantly reduced alcohol lever-responding in both non-dependent and dependent rat groups (****p<0.0001; **p<0.005, Tukey’s post hoc test) (Fig 8A). Consistently, alcohol dependent rats self-administered greater amounts of alcohol than non-dependent rats (1.0 *vs* 0.6 g/kg/30-min session; main effect of group: F (1,103) = 21.85; p<0.0001). Two-way ANOVA revealed a significant main effect of CYM5442 treatment on alcohol intake (F (3,103) = 18.73; p<0.0001) with no significant interaction between group and treatment (F (3,103) = 1.553; p>0.05). Post-hoc comparisons revealed that only 10 mg/kg dose of CYM5442 significantly decreased alcohol self-administration in both non-dependent and dependent rat groups (****p<0.0001; **p<0.005, by Tukey’s post hoc test) (Fig 8B).

**Fig 8.**
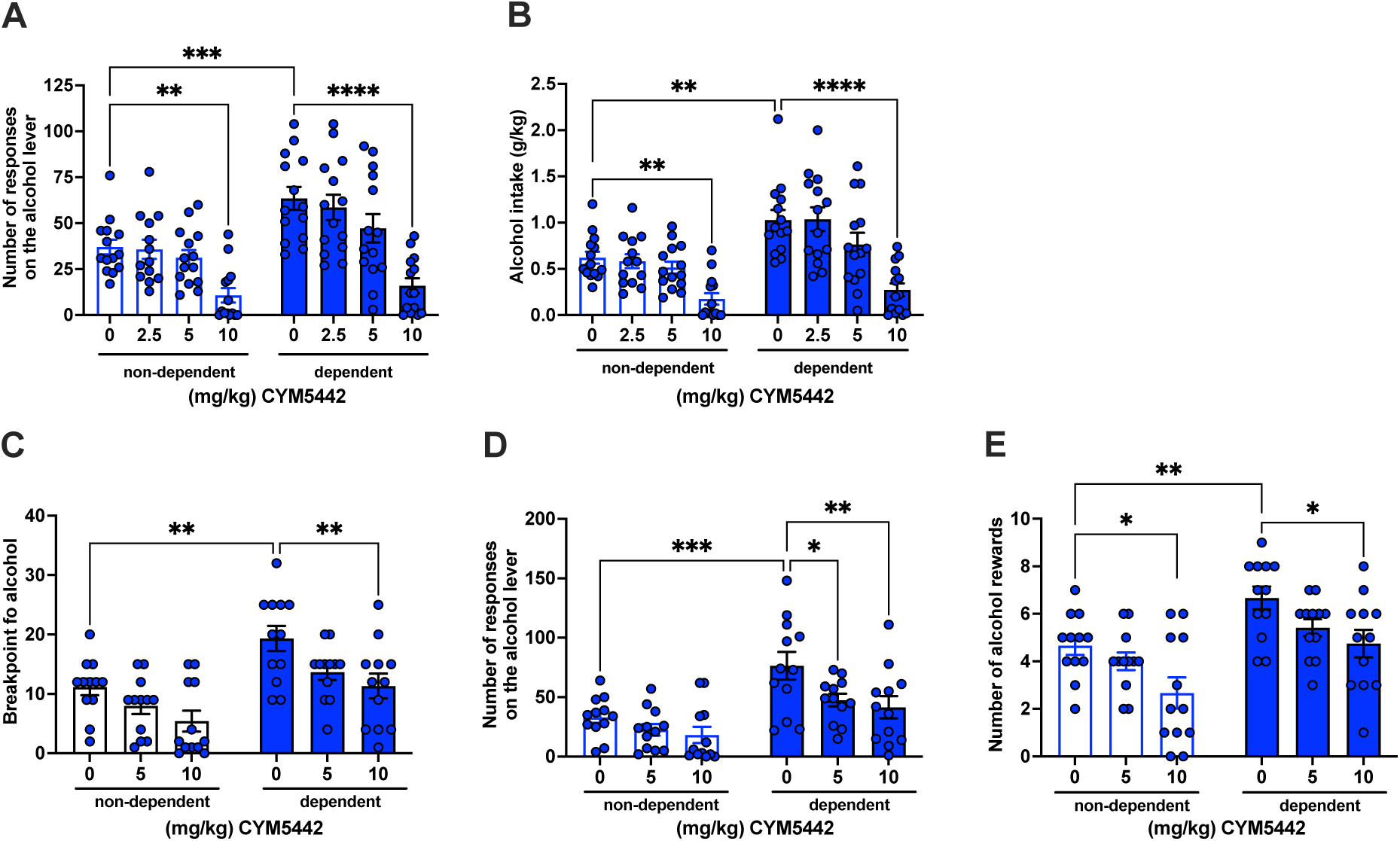
The selective S1P receptor 1 agonist CYM5442 reduces oral alcohol self-administration under fixed and progressive ratio schedules of reinforcement in both non-dependent and dependent male rats. **A)** Number of responses on the alcohol lever and **B)** amount of self-administered alcohol were reduced by administration of CYM5442, 60 min prior to the start of the 30-min drinking session. (**p< 0.005; ***p<0.001; ****p<0.0001). Each bar represents the mean ± SEM of n=14-15 rats. Similarly, Breakpoint for alcohol **(C)**, number of responding on the alcohol lever **(D)** and number of alcohol rewards **(E)** were reduced by administration of CYM5442, 60 min prior to the start of the 60-min drinking session (*p<0.05; **p< 0.005; ***p<0.0005). Each bar represents the mean ± SEM of n=12 rats.

To determine whether CYM5442 also affected the motivational properties of alcohol, non-dependent and dependent rats were exposed to a progressive ratio schedule of reinforcement. We observed that alcohol dependent rats exhibited a significantly higher motivation to obtain alcohol compared to non-dependent rats, as indicated by higher break point (BP) value. (19.3 vs 11.7). Two-way ANOVA revealed significant main effects of both group (F (1,66) = 22.13; p<0.0001)) and treatment (F (2,66) = 8.26; p<0.001) with no significant interaction between the two factors (F (2,66) = 0.32; p>0.05). Post-hoc analysis showed that only the 10 mg/kg dose of CYM5442 significantly reduced the motivation for alcohol in both non-dependent and dependent rat groups (**p <0.005; *p=0.05 by Tukey’s post hoc test) (Fig 8C). Consistently, the number of responses on the active lever was significantly higher in the alcohol dependent rat group than the non-dependent rat group (76.3 vs 33.1 responses over 60-min session). Two-way ANOVA revealed significant main effects of group (F (1,66) = 24.04; p<0.0001) and treatment (F (2,66) = 5.98; p<0.005), but no significant interaction (F (2,66) =1.06; p>0.05). Post-hoc analysis revealed that both 5 and 10 mg/kg doses of CYM5442 significantly reduced lever-responding for alcohol in the dependent rat group only (**p<0.005; *p<0.05 by Tukey’s post hoc test) (Fig 8D). Similar results were observed on number of alcohol rewards earned. Two-way ANOVA revealed significant main effects of group (F (1,66) = 21.33; p<0.0001) and treatment (F (2,66) = 8.11; p<0.005), but no significant interaction (F (2,66) = 0.28; p>0.05). Post-hoc analysis revealed that 10 mg/kg dose of CYM5442 significantly reduced lever-responding for alcohol in both the non-dependent and dependent rat groups (**p<0.005; *p<0.05 by Tukey’s post hoc test) (Fig 8E).

#### CYM5442 prevents cues-induced reinstatement of alcohol seeking behavior in alcohol dependent rats

After the PR test, rats completed 10 regular 30-min sessions of alcohol self-administration and then underwent extinction responding training. Alcohol dependent and non-dependent rats progressively extinguished their alcohol seeking behavior over five consecutive days. Two-way ANOVA revealed significant main effects of group (F (1,62)=25.86; p<0.0001) and day [F (4,247) =22.07; p<0.0001) but no significant interaction between these factors (F (4,247) =1.53; p>0.05). Post hoc analysis indicated that lever-responding was higher in dependent rats compared to non-dependent rats during extinction days 1-3 (****p<0.0001, ***p<0.0005, *p<0.05 by Tukey’s post hoc test) (Fig 9A). On the day following the last extinction session, dependent and non-dependent rats were allocated into two groups based on their number of responses on the active lever during the last two days of extinction. (Fig 9B, left panel). Rats of both groups received either vehicle (0 mg/kg) or CYM5442 (10 mg/kg) 60-min before the reinstalment session. Three-way ANOVA revealed a significant main effect of group (F (1,60) = 8.48; p<0.005) and treatment (F (1,60) = 8.64; p<0.005) but not protocol (extinction/reinstatement) (F (1,60) = 0.847; p>0.05). No significant interactions were observed for protocol x treatment (F (1,60) = 0.04; p>0.05), protocol x group (F (1,60) = 21.57; p>0.05), treatment x group (F (1,60) = 0.949; p>0.05), or protocol x group x treatment (F (1,60) = 2.38; p>0.05). Post hoc analysis revealed that CYM5442 significantly prevented cue-induced reinstatement in the dependent rat group ($p<0.005; #p<0.05; ***p<0.0001, by Bonferroni post hoc). (Fig 9B, right panel).

**Fig 9.**
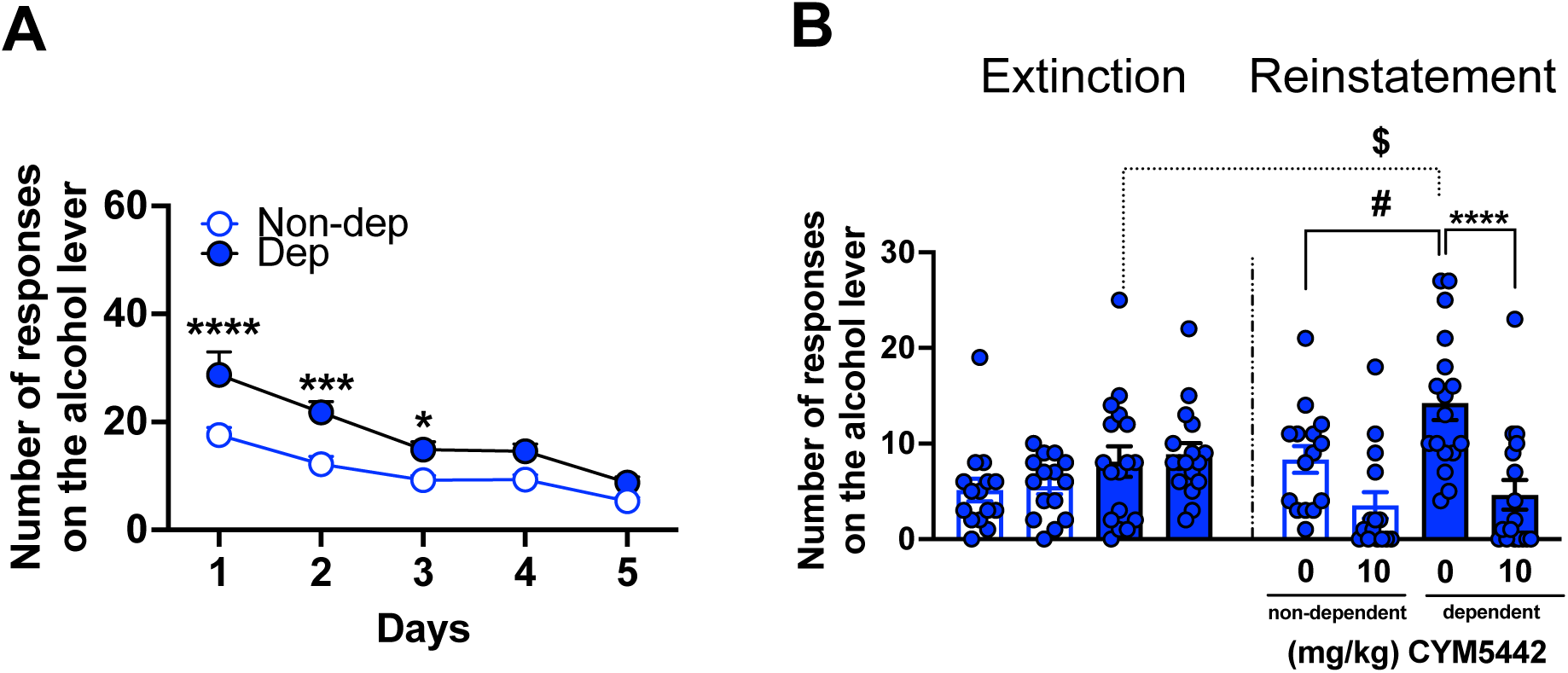
The selective S1P receptor 1 agonist CYM5442 prevents cues-induced reinstatement of alcohol seeking behavior in non-dependent and dependent rats. After the PR test, rats were re-exposed to ten 30-min alcohol self-administration sessions until lever-responding stabilized. Subsequently, rats underwent an extinction-responding training made up of 5 consecutive 30-min daily sessions during which lever-responding was unreinforced, and pumps, stimulus light, and tone generator were off. **A)** Over the 5 sessions, rats progressively reduced their lever-responding behavior (*p<0.05; ***p<0.0005; ****p< 0.0001). Each point represents the mean ± SEM of 28 (non-dependent) and 34 (dependent) rats. **B)** Rats’ alcohol-seeking behavior was reinstated by the repeated presentation of a complex of visual and auditory stimuli previously associated with alcohol availability. Administration of CYM5442 60 min before the start of the 30-min test prevented cues-induced reinstatement in dependent rats. (3-way ANOVA followed by Šidák post hoc test; #p< 0.05; $p< 0.005; ****p<0.0001). Each bar represents the mean ± SEM of n=14 (non-dependent) and 17 (dependent) rats.

#### CYM5442 does not alter rat spontaneous locomotor activity

As for mice, potential secondary effects of CYM5442 were assessed on spontaneous locomotor activity of alcohol-naive rats. One-way ANOVA analyzing the total ambulatory distance traveled by rats during a 60-minute session showed that administration of CYM5442 did not significantly affect locomotor activity (F (3,20) = 0.59; p>0.05) (Fig 10A). Furthermore, two-way RM ANOVA of ambulatory distance across six 10-min intervals, revealed a significant effect of time (F(5,100) = 156.7; p<0.0001), but no significant effects of treatment (F (3,20) = 0.59; p> 0.05) or time x treatment (F (15,100) = 0.71; p>0.05) (Fig. 10B).

**Fig 10.**
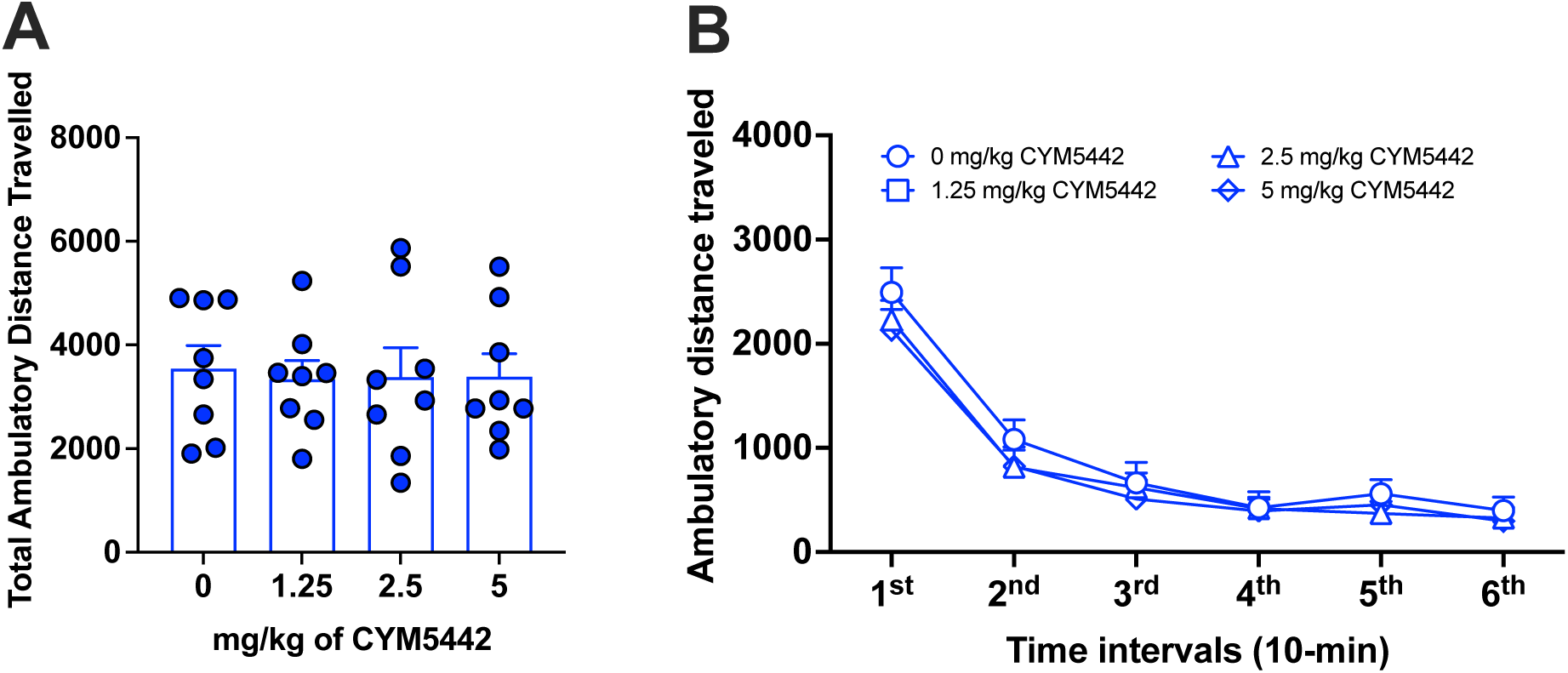
The selective S1P receptor 1 agonist CYM5442 does not alter spontaneous locomotor activity of alcohol-naive rats. **A)** Total ambulatory distance travelled, **B)** Ambulatory distance travelled over the six 10-min intervals. CYM5442 was administered IP 60 min before the start of the 1 h locomotor activity test. Each bar represents the mean ± SEM of n=6 rats.

#### S1P regulates a complex set of genes in the transition to alcohol dependence

We carried out RNA-Seq of the PFC of CYM5442 and vehicle treated rats with histories of CIE and alcohol-naive controls, Fig. 11. Pathway analysis by gene set enrichment analysis (GSEA) revealed that treatment with CYM5442 affect several several key pathways related to signal transduction, neuronal function, synaptic and structural neuronal plasticity, and regulation of gene expression. These results suggest that S1P1-regulated pathways are complex gene expression program with the potential to substantially affect neuronal states consistent with the role of S1P1 in the transition to escalated (dependent) alcohol intake, Fig. 11.

**Fig. 11.**
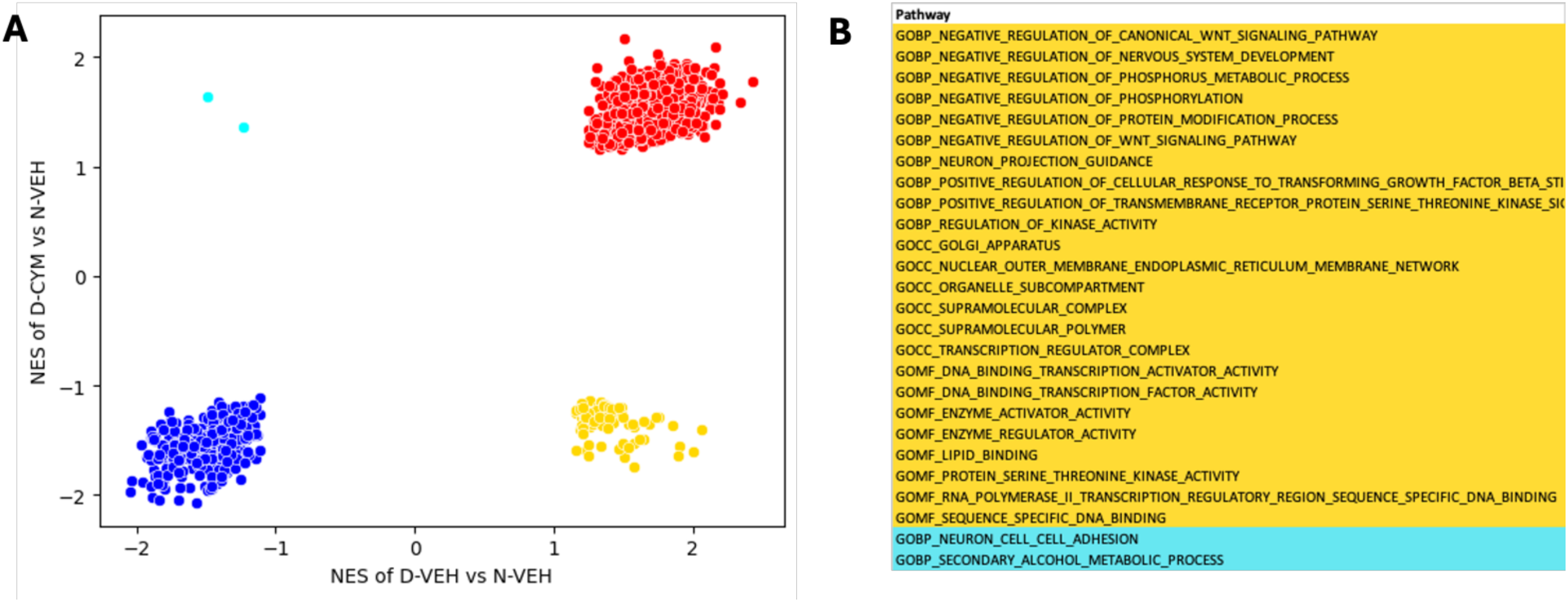
Pathway analysis of CYM5442-treated rats exposed to CIE. **A)** The GSEA algorithm was used for pathway analysis of gene expression changes induced by CYM5442 in rats exposed to CIE vs. vehicle-treated rats exposed to CIE. On the X axis is the pathway analysis of CIE-exposed (dependent) rats vs. alcohol naive control rats (D-VEH v N-VEH); on the Y axis is the pathway analysis of CIE-exposed CYM5442-treated rats vs. alcohol naive control rats (D-CYM v N-VEH). Concordantly regulated pathways in vehicle-treated CIE-exposed rats vs. controls and CYM5442-treated CIE-exposed rats vs. controls are shown on the top right (red: induced) and bottom left (blue: decreased). On the top left (light green) and bottom right (yellow) are the pathways discordantly regulated by vapor in CYM5442 treated vs. vehicle treated rats. The normalized enrichment scores (NES) are shown for pathways that are significantly differential regulated (p<0.05). **B)** Representative CYM5442-regulated pathways. The pathways affected by CYM5442 treatment in CIE exposed rats included pathways representative of signal transduction, neuronal function, synaptic and structural neuronal plasticity, and regulation of gene expression.

## Discussion

Here we show that S1P levels were decreased after administration of an intoxicating dose of alcohol in the mouse prefrontal cortex (PFC), a key brain region in the shift from moderate to compulsive alcohol drinking and taking (19–28). Also relevantly, ceramide, the precursor of sphingosine, was reduced in the forebrain of male selectively bred alcohol-preferring rats with a history of chronic intermittent drinking of 20% alcohol (29).

High levels of S1P_1_ receptor have been identified in the prefrontal cortex and striatum, two brain regions involved in alcohol use and abuse (30). In human alcoholics, prefrontal cortex deficits are believed to contribute to excessive drinking and increased vulnerability to relapse (21–28, 31). Rodents with a history of alcohol dependence exhibit cognitive impairment that reflects prefrontal cortex dysfunction (25). Fronto-striatal circuits are implicated in the loss of control and enhanced motivation to drink that characterize AUD (32–35).

Acute administration of CYM5442 effectively prevented the consumption of intoxicating amounts of alcohol in male and female C57BL/6J mice exposed to the DID paradigm. Alcohol intake was reduced in a dose-dependent manner, with a significant effect observed at both 2.5 and 5 mg/kg CYM5442. Notably, the effect of 5 mg/kg CYM5442 persisted throughout the 4 h drinking session in both sexes despite its short half-life due to rapid clearance from circulation (16). Furthermore, the highest dose of CYM5442 (5 mg/kg) also reduced alcohol intake in both non-dependent and dependent male and female C57BL/6J mice in the 2-bottle choice paradigm after chronic intermittent exposure to alcohol vapor. CYM5442 did not alter the hypnotic/sedative effects of alcohol as assessed by the loss of righting reflex or interfere with its metabolism in male and female C57BL/6J mice. Notably, in the conditioned place aversion test, CYM5442 was less aversive than naltrexone, an FDA-approved medication for AUD in humans.

Parallel studies in male Wistar rats confirmed the ability of CYM5442 to interfere with the reinforcing and motivational properties of alcohol as evidenced by reduced self-administration under both fixed and progressive ratio schedule of reinforcement in non-dependent and dependent rats. Additionally, in dependent rats, CYM5442 prevented cues-induced reinstatement of alcohol seeking behavior, a validated model of loss of control over alcohol and relapse into heavy alcohol drinking (36).

To evaluate CYM5442 specificity, we tested CYM5442 effects on saccharin and sucrose intake, two palatable reinforcers with different caloric value, using an experimental design paralleling the DID paradigm, with independent groups of male and female mice. CYM5442 significantly reduced saccharin intake in both sexes, but only at a higher dose than that effective in reducing alcohol intake. In contrast, sucrose intake was reduced in males at 2.5 and 5 mg/kg whereas in females only at 5 mg/kg, with no effect detected at the end of the 4 h drinking session. Additionally, CLAMS data indicate that acute CYM5442 also affects food, water intake, and respiratory exchange ratio. Importantly, CYM5442 at doses between 1.25 - 5 mg/kg did not alter mouse motor coordination in the rotarod apparatus, ruling out nonspecific effects such as sedation or malaise as explanations for reduced drinking and feeding.

Overall, these results indicating CYM5442 effects on mouse drinking (drug, non-drug reinforcers, and water) and feeding, suggest that the S1P_1_ receptor could be also involved in general consummatory behavior. That is akin to other drugs that reduce alcohol intake. Importantly, naltrexone, an FDA-approved medication for the treatment of AUD, reduces alcohol, sucrose, and saccharin intake of male C57BL/6J mice exposed to the DID paradigm (37). However, in contrast with the observed CYM5442 properties, naltrexone also induces conditioned place aversion (38). A similar ability to effect broad reduction of rodent consummatory behavior has been shown for cannabinoids 1 (CB_1_) receptors (CB_1_-R) antagonists such as rimonabant, AM6527, and AM4113 (39, 40). Interestingly, it has been reported that S1P and its analog, the non-selective S1P agonist fingolimod, interact with the CB_1_-Rs, suggesting that molecules belonging to the same pharmacological class could potentially be provided with CB_1_-R activity (41). However, we found that CYM5442 does not (see Supplementary material), supporting that the effect of S1P agonists on alcohol drinking is independent of CB_1_-R. Other drugs that cause broad inhibition of consummatory behaviors include the CRF1-selective antagonist NBI-27914 and the immune-targeting compound tacrolimus that reduced both alcohol and saccharin intake in the DID paradigm (42). The glucagon-like peptide-1 (GLP-1) analogue semaglutide has been shown to reduce alcohol, water, saccharin, maltodextrin and corn oil intake (43). Overall, the present findings are consistent with the overlapping neurobiological and chemosensory mechanisms that regulate food, drug reward, and consummatory behavior (44). Consistently, preclinical and clinical data indicate a strong correlation between excessive alcohol and sweet food consumption in rodents and humans (45–47) and drug abuse and binge eating disorders have been shown to share imbalances in brain systems that regulate motivation, reward saliency, decision-making, and self-control (44, 48, 49).

In the present study, we observed that acute administration of CYM5442 effectively reduced food intake and RER in mice, supporting a role for S1P_1_ also in energy balance and metabolism. This effect could be explained by the activation of S1P_1_ receptors localized at the level of the hypothalamus, the regulatory center of feeding and drinking. In this regard, recent studies have shown high levels of S1P_1_ protein in all hypothalamic nuclei with a predominance in the arcuate, dorsomedial, and ventromedial nuclei. In addition, the S1P_1_ receptor was found predominantly in anorexigenic but not in orexigenic neurons in the arcuate nucleus (50, 51).

In conclusion, we showed that the selective S1P receptor agonists reduced alcohol intake and consummatory behavior, particularly on caloric reinforcers with low aversion potential. These results establish S1P signaling as a therapeutic target for AUD.

## Conflict of Interest

No conflict of interest to declare.

## Funding

This work was supported by NIH Grant AA021667; IL was partially supported by training grant T32 AA007456.

**All authors approved the final version of the manuscript**

## Notes

### Competing Interest Statement

The authors have declared no competing interest.

## References

1. Litten RZ, Ryan ML, Falk DE, Reilly M, Fertig JB, Koob GF. 2015. Heterogeneity of alcohol use disorder: understanding mechanisms to advance personalized treatment. Alcohol Clin Exp Res 39:579–84.

2. Litten RZ, Wilford BB, Falk DE, Ryan ML, Fertig JB. 2016. Potential medications for the treatment of alcohol use disorder: An evaluation of clinical efficacy and safety. Subst Abus 37:286–98.

3. Litten RZ, Falk DE, Ryan ML, Fertig J, Leggio L. 2020. Five Priority Areas for Improving Medications Development for Alcohol Use Disorder and Promoting Their Routine Use in Clinical Practice. Alcohol Clin Exp Res 44:23–35.

4. Karunakaran I, van Echten-Deckert G. 2017. Sphingosine 1-phosphate - A double edged sword in the brain. Biochim Biophys Acta Biomembr 1859:1573–1582.

5. Blondeau N, Lai Y, Tyndall S, Popolo M, Topalkara K, Pru JK, Zhang L, Kim H, Liao JK, Ding K, Waeber C. 2007. Distribution of sphingosine kinase activity and mRNA in rodent brain. J Neurochem 103:509–17.

6. Fukuda Y, Kihara A, Igarashi Y. 2003. Distribution of sphingosine kinase activity in mouse tissues: contribution of SPHK1. Biochem Biophys Res Commun 309:155–60.

7. Blaho VA, Hla T. 2014. An update on the biology of sphingosine 1-phosphate receptors. J Lipid Res 55:1596–608.

8. Novgorodov AS, El-Alwani M, Bielawski J, Obeid LM, Gudz TI. 2007. Activation of sphingosine-1-phosphate receptor S1P5 inhibits oligodendrocyte progenitor migration. FASEB J 21:1503–14.

9. Dusaban SS, Chun J, Rosen H, Purcell NH, Brown JH. 2017. Sphingosine 1-phosphate receptor 3 and RhoA signaling mediate inflammatory gene expression in astrocytes. J Neuroinflammation 14:111.

10. Gril B, Paranjape AN, Woditschka S, Hua E, Dolan EL, Hanson J, Wu X, Kloc W, Izycka-Swieszewska E, Duchnowska R, Peksa R, Biernat W, Jassem J, Nayyar N, Brastianos PK, Hall OM, Peer CJ, Figg WD, Pauly GT, Robinson C, Difilippantonio S, Bialecki E, Metellus P, Schneider JP, Steeg PS. 2018. Reactive astrocytic S1P3 signaling modulates the blood-tumor barrier in brain metastases. Nat Commun 9:2705.

11. Lee JY, Han SH, Park MH, Baek B, Song IS, Choi MK, Takuwa Y, Ryu H, Kim SH, He X, Schuchman EH, Bae JS, Jin HK. 2018. Neuronal SphK1 acetylates COX2 and contributes to pathogenesis in a model of Alzheimer’s Disease. Nat Commun 9:1479.

12. Cahalan SM, Gonzalez-Cabrera PJ, Nguyen N, Guerrero M, Cisar EA, Leaf NB, Brown SJ, Roberts E, Rosen H. 2013. Sphingosine 1-phosphate receptor 1 (S1P(1)) upregulation and amelioration of experimental autoimmune encephalomyelitis by an S1P(1) antagonist. Mol Pharmacol 83:316–21.

13. Proia RL, Hla T. 2015. Emerging biology of sphingosine-1-phosphate: its role in pathogenesis and therapy. J Clin Invest 125:1379–87.

14. McGinley MP, Cohen JA. 2021. Sphingosine 1-phosphate receptor modulators in multiple sclerosis and other conditions. Lancet 398:1184–1194.

15. Rhodes JS, Best K, Belknap JK, Finn DA, Crabbe JC. 2005. Evaluation of a simple model of ethanol drinking to intoxication in C57BL/6J mice. Physiol Behav 84:53–63.

16. Gonzalez-Cabrera PJ, Jo E, Sanna MG, Brown S, Leaf N, Marsolais D, Schaeffer MT, Chapman J, Cameron M, Guerrero M, Roberts E, Rosen H. 2008. Full pharmacological efficacy of a novel S1P1 agonist that does not require S1P-like headgroup interactions. Mol Pharmacol 74:1308–18.

17. Becker HC, Lopez MF. 2004. Increased ethanol drinking after repeated chronic ethanol exposure and withdrawal experience in C57BL/6 mice. Alcohol Clin Exp Res 28:1829–38.

18. Finn DA, Snelling C, Fretwell AM, Tanchuck MA, Underwood L, Cole M, Crabbe JC, Roberts AJ. 2007. Increased drinking during withdrawal from intermittent ethanol exposure is blocked by the CRF receptor antagonist D-Phe-CRF(12-41). Alcohol Clin Exp Res 31:939–49.

19. Goldstein RZ, Leskovjan AC, Hoff AL, Hitzemann R, Bashan F, Khalsa SS, Wang GJ, Fowler JS, Volkow ND. 2004. Severity of neuropsychological impairment in cocaine and alcohol addiction: association with metabolism in the prefrontal cortex. Neuropsychologia 42:1447–58.

20. Bergman H, Engelbrektson K, Fransson A, Herlitz K, Hindmarsh T, Neiman J. 1998. Alcohol-induced cognitive impairment is reversible. Neuropsychological tests but not MRT show improvement after abstinence. Lakartidningen 95:4228, 4231–6.

21. Rando K, Hong KI, Bhagwagar Z, Li CS, Bergquist K, Guarnaccia J, Sinha R. 2011. Association of frontal and posterior cortical gray matter volume with time to alcohol relapse: a prospective study. Am J Psychiatry 168:183–92.

22. Sabia S, Gueguen A, Berr C, Berkman L, Ankri J, Goldberg M, Zins M, Singh-Manoux A. 2011. High alcohol consumption in middle-aged adults is associated with poorer cognitive performance only in the low socio-economic group. Results from the GAZEL cohort study. Addiction 106:93–101.

23. Sullivan EV, Rosenbloom MJ, Pfefferbaum A. 2000. Pattern of motor and cognitive deficits in detoxified alcoholic men. Alcohol Clin Exp Res 24:611–21.

24. Tedstone D, Coyle K. 2004. Cognitive impairments in sober alcoholics: performance on selective and divided attention tasks. Drug Alcohol Depend 75:277–86.

25. Trantham-Davidson H, Burnett EJ, Gass JT, Lopez MF, Mulholland PJ, Centanni SW, Floresco SB, Chandler LJ. 2014. Chronic alcohol disrupts dopamine receptor activity and the cognitive function of the medial prefrontal cortex. J Neurosci 34:3706–18.

26. Wolwer W, Burtscheidt W, Redner C, Schwarz R, Gaebel W. 2001. Out-patient behaviour therapy in alcoholism: impact of personality disorders and cognitive impairments. Acta Psychiatr Scand 103:30–7.

27. Finn PR, Justus A, Mazas C, Steinmetz JE. 1999. Working memory, executive processes and the effects of alcohol on Go/No-Go learning: testing a model of behavioral regulation and impulsivity. Psychopharmacology (Berl) 146:465–72.

28. Koob GF, Volkow ND. 2010. Neurocircuitry of addiction. Neuropsychopharmacology 35:217–38.

29. Godfrey J, Jeanguenin L, Castro N, Olney JJ, Dudley J, Pipkin J, Walls SM, Wang W, Herr DR, Harris GL, Brasser SM. 2015. Chronic Voluntary Ethanol Consumption Induces Favorable Ceramide Profiles in Selectively Bred Alcohol-Preferring (P) Rats. PLoS One 10:e0139012.

30. Jiang H, Joshi S, Liu H, Mansor S, Qiu L, Zhao H, Whitehead T, Gropler RJ, Wu GF, Cross AH, Benzinger TLS, Shoghi KI, Perlmutter JS, Tu Z. 2021. In Vitro and In Vivo Investigation of S1PR1 Expression in the Central Nervous System Using [(3)H]CS1P1 and [(11)C]CS1P1. ACS Chem Neurosci 12:3733–3744.

31. Bergman H, Engelbrektson K, Fransson A, Herlitz K, Hindmarsh T, Neiman J. 1998. [Alcohol-induced cognitive impairment is reversible. Neuropsychological tests but not MRT show improvement after abstinence]. Lakartidningen 95:4228, 4231–6.

32. Volkow ND, Wiers CE, Shokri-Kojori E, Tomasi D, Wang GJ, Baler R. 2017. Neurochemical and metabolic effects of acute and chronic alcohol in the human brain: Studies with positron emission tomography. Neuropharmacology 122:175–188.

33. Jeanblanc J, He DY, Carnicella S, Kharazia V, Janak PH, Ron D. 2009. Endogenous BDNF in the dorsolateral striatum gates alcohol drinking. J Neurosci 29:13494–502.

34. Corbit LH, Nie H, Janak PH. 2014. Habitual responding for alcohol depends upon both AMPA and D2 receptor signaling in the dorsolateral striatum. Front Behav Neurosci 8:301.

35. Chen J, Nam HW, Lee MR, Hinton DJ, Choi S, Kim T, Kawamura T, Janak PH, Choi DS. 2010. Altered glutamatergic neurotransmission in the striatum regulates ethanol sensitivity and intake in mice lacking ENT1. Behav Brain Res 208:636–42.

36. Martin-Fardon R, Weiss F. 2013. Modeling relapse in animals. Curr Top Behav Neurosci 13:403–32.

37. Morales I, Rodriguez-Borillo O, Font L, Pastor R. 2020. Effects of naltrexone on alcohol, sucrose, and saccharin binge-like drinking in C57BL/6J mice: a study with a multiple bottle choice procedure. Behav Pharmacol 31:256–271.

38. Kuzmin A, Sandin J, Terenius L, Ogren SO. 2003. Acquisition, expression, and reinstatement of ethanol-induced conditioned place preference in mice: effects of opioid receptor-like 1 receptor agonists and naloxone. J Pharmacol Exp Ther 304:310–8.

39. Sink KS, Vemuri VK, Wood J, Makriyannis A, Salamone JD. 2009. Oral bioavailability of the novel cannabinoid CB1 antagonist AM6527: effects on food-reinforced behavior and comparisons with AM4113. Pharmacol Biochem Behav 91:303–6.

40. Aravamudan VM, Er C, Hussain I, Cheong NWW, Chern Hao C, Kuthah N, En-Xian Tan E, Fan BE. 2018. A Case of Parvovirus-Related Haemophagocytic Lymphohistiocytosis in a Patient with HbH Disease. Case Rep Med 2018:8057045.

41. Paugh SW, Cassidy MP, He H, Milstien S, Sim-Selley LJ, Spiegel S, Selley DE. 2006. Sphingosine and its analog, the immunosuppressant 2-amino-2-(2-[4-octylphenyl]ethyl)-1,3-propanediol, interact with the CB1 cannabinoid receptor. Mol Pharmacol 70:41–50.

42. Grigsby KB, Savarese AM, Metten P, Mason BJ, Blednov YA, Crabbe JC, Ozburn AR. 2020. Effects of Tacrolimus and Other Immune Targeting Compounds on Binge-Like Ethanol Drinking in High Drinking in the Dark Mice. Neurosci Insights 15:2633105520975412.

43. Chuong V, Farokhnia M, Khom S, Pince CL, Elvig SK, Vlkolinsky R, Marchette RC, Koob GF, Roberto M, Vendruscolo LF, Leggio L. 2023. The glucagon-like peptide-1 (GLP-1) analogue semaglutide reduces alcohol drinking and modulates central GABA neurotransmission. JCI Insight 8.

44. Volkow ND, Wang GJ, Tomasi D, Baler RD. 2013. Obesity and addiction: neurobiological overlaps. Obes Rev 14:2–18.

45. Kampov-Polevoy A, Garbutt JC, Janowsky D. 1997. Evidence of preference for a high-concentration sucrose solution in alcoholic men. Am J Psychiatry 154:269–70.

46. Kampov-Polevoy AB, Garbutt JC, Janowsky DS. 1999. Association between preference for sweets and excessive alcohol intake: a review of animal and human studies. Alcohol Alcohol 34:386–95.

47. Leggio L, Addolorato G, Cippitelli A, Jerlhag E, Kampov-Polevoy AB, Swift RM. 2011. Role of feeding-related pathways in alcohol dependence: A focus on sweet preference, NPY, and ghrelin. Alcohol Clin Exp Res 35:194–202.

48. Volkow ND, Wise RA, Baler R. 2017. The dopamine motive system: implications for drug and food addiction. Nat Rev Neurosci 18:741–752.

49. Wiss DA, Avena N, Rada P. 2018. Sugar Addiction: From Evolution to Revolution. Front Psychiatry 9:545.

50. Silva VR, Katashima CK, Bueno Silva CG, Lenhare L, Micheletti TO, Camargo RL, Ghezzi AC, Camargo JA, Assis AM, Tobar N, Morari J, Razolli DS, Moura LP, Pauli JR, Cintra DE, Velloso LA, Saad MJ, Ropelle ER. 2016. Hypothalamic S1P/S1PR1 axis controls energy homeostasis in Middle-Aged Rodents: the reversal effects of physical exercise. Aging (Albany NY) 9:142–155.

51. Silva VR, Micheletti TO, Pimentel GD, Katashima CK, Lenhare L, Morari J, Mendes MC, Razolli DS, Rocha GZ, de Souza CT, Ryu D, Prada PO, Velloso LA, Carvalheira JB, Pauli JR, Cintra DE, Ropelle ER. 2014. Hypothalamic S1P/S1PR1 axis controls energy homeostasis. Nat Commun 5:4859.

