## Supplementary material for "Sphingosine-1-phosphate (S1P) signaling as a novel therapeutic target for alcohol abuse": Suppl Material

**Table of Contents:**

1. **Experimental procedures**
2. **Total RNA Isolation and RNA-Sequencing**.
3. **Supplemental Fig. 1: Lack of CB1 activity of CYM-5442.**
4. **References**

1. **Experimental procedures.**

**Mouse experiments**

*Targeted LC-MS/MS quantification of sphingosine-1-phosphate in the mouse brain*

Tissue was extracted with glass beads and 600uL of cold MeOH: H2O (4:1) added. Tissue samples were homogenized with glass beads in homogenizer, and sonicated in ice bath for 10 min. The homogenized solution was then transferred to 1.5 mL Eppendorf vial, rinsed the glass vial with additional 200 μL cold MeOH: H2O (4:1, v/v) and combine the wash solution to the same Eppendorf vial. Internal standard added during extraction. To precipitate proteins, the samples was incubated for 1 hr at −20 °C then followed by 15 min centrifugation at 13,000 rpm and 4 °C. The supernatant was transferred then evaporated to dryness in a vacuum concentrator. Dry extracts were resuspended in 100uL MeOH, centrifuged 15 min at 13000 rpm and 4 °C to remove insoluble debris and supernatant transferred to autosampler vials for analysis. The samples were run on an Agilent 6495 triple quadrupole mass spectrometer coupled to an Agilent 1290 UPLC stack. Column used was an Agilent poroshell 2.1x50mm C18. The mobile phase was composed of A= Water/0.1% formic acid, and B= ACN/0.1% formic acid. The flowrate was set to 300μL/min, and the sample injection volume was 5 μL. Gradient was: T0 90/10, T5 5/95 T8 off. There was a 3.5-minute re-equilibration step. Operating in positive ion mode, the source conditions were as follows: Capillary voltage = 3500V, Gas temp = 120C, Gas flow = 13L/min, Nebulizer pressure = 25psi. Sheath gas temp = 350C, Sheath gas flow = 12L/min. Nozzle voltage was set to 2000V. Data was processed using Agilent Quantitative analysis software. S1P d17 was added as an internal standard at 500nM. Calibration standards were run from 50nM to 10uM. Results are given as fmol/mg.

*Drinking the Dark (DID) paradigm*

The acute effect of the S1P agonists was assessed using a binge-like alcohol drinking model in C57BL/6J mice employing the DID paradigm. As described by Rhodes and colleagues (1), three hours after lights-off, the water bottle of single-housed mice was replaced with a bottle containing 20% (v/v) alcohol and left in place for 2 hours. This procedure was repeated over three consecutive days to allow stabilization of mice alcohol intake. On day 4, mice received an IP injection of either fingolimod, ozanimod or CYM5442 before the drinking session, which had a duration of 4 h. Alcohol intake was measured after the first and second 2 h interval. The amount of alcohol consumed was determined by weighing the bottles (± 0.01 g accuracy) immediately before and after each drinking session and expressed as grams of pure alcohol per kilogram of body weight (g/kg).

*Non-drug reinforces in the Drinking in the Dark paradigm*

To assess effects on non-drug reinforces, separate cohorts of male and female mice underwent the same DID schedule with saccharin (0.002% w/v) or sucrose (0.05% w/v) solutions. On day 4, mice received an IP injection of CYM5442 or vehicle before a 4-h session. Fluid intake was measured after the first 2 h and at the end of the session, expressed as ml/30 g body weight.

*2-bottle choice paradigm and Chronic intermittent access (CIE) to alcohol vapors*

Male and female C57BL/6J mice were singly housed and exposed to a 2-h, 2-bottle choice (15% alcohol vs. water) paradigm for 5 consecutive days, with consumption measured by weighing the bottles before and after the drinking session. Mice were then divided into two balanced groups based on baseline intake: one (dependent) received intermittent ethanol vapor (CIE; 16 h ON/9 h OFF) and the other (non-dependent) received air. Before vapor exposure, dependent mice were injected IP with 1.5 g/kg alcohol (15% w/v) plus 68.1 mg/kg pyrazole and placed in vapor chambers (La Jolla Ethanol Research, CA, USA). Tail blood was collected daily to monitor BALs (target 150–200 mg%). Seventy-two hours after vapor removal, mice had 2-h access to 15% alcohol vs. water for 5 consecutive days. This cycle of vapor/air exposure followed by 2BC testing was repeated six times.

*Food, water intake, and RER*

The effect of CYM5442 was evaluated on food (g), water intake (ml), and respiratory exchange ratio (RER: VCO2/VO2) of male C57BL/6J mice allocated into CLAMS (Comprehensive Lab Animal Monitoring System, Columbus Instruments, Columbus, OH USA). After 1-week acclimatization, on test day, mice were removed from metabolic chambers, injected with CYM5442 60 min before 3 h into the dark of the light/dark cycle, and then placed back into chambers.

*Conditioned Place Aversion.*

Male C57BL/6J mice were tested in the conditioned place aversion paradigm. The apparatus consisted of two compartments, separated by a guillotine door, provided with different tactile stimuli (MedAssociate, VT, USA). Experimental protocol was divided into three phases: I) Spontaneous choice assessment (days 1-5): each mouse was positioned in the apparatus and left free to explore it for 15 min, with the guillotine door open, to assess their spontaneous preference between the two compartments. For each mouse, pre-conditioning time was calculated as the average of the time spent in the preferred compartment for the last 3 days of spontaneous preference. II) Conditioning (days 6-15): mice were divided into three balanced groups based on their pre-conditioning time and IP injected with either vehicle, naltrexone (NTX, 16 mg/kg), or CYM5442 (5 mg/kg). Subsequently, they were immediately confined in the preferred compartment for 1h, with the guillotine door closed. Mice that were confined in the preferred compartment one day were treated with vehicle and confined in the other compartment the day after. Hence, mice were conditioned on alternative days for 10 days. III) Assessment of shifted spontaneous preference (day 16) drug-free mice were exposed to the apparatus and left free to explore it for 15 min, with the guillotine door open, to assess whether their spontaneous preference changed and to determine the post-conditioning time. Difference between pre-and post-conditioning time was measured.

*Loss of Righting Reflex and alcohol metabolism*

Male and female alcohol-naive C57BL/6J mice were administered with either saline or CYM5442 (2.5 mg/kg) 60 min before receiving an IP injection of alcohol 15% (w/v) (3.5 g/kg). Immediately after, when the mouse showed the first signs of sedation, it was placed on its back on a V-shaped position into a cleaned cage, left undisturbed and monitored. Duration of loss of righting reflex (LORR) was measured, and blood samples were collected from the tip of the tail of each mouse.

*Rotarod test*

Motor coordination was assessed in male and female C57BL/6J mice in the rotarod apparatus (Rotamex-5, Columbus Instruments, USA). Mice were habituated to handling and IP injections the week before testing. On test day, mice were injected with either CYM5442 or vehicle 60 min before the first trial. Each mouse underwent 3 trials of 300 s (intertrial interval 60 s) with an acceleration from 4 to 40 rpm; trials in which animals fell before 5 s were repeated. Latency to fall was expressed as the average values over the 3 trials.

**Rat experiments.**

*Alcohol self-administration*

*Apparatus*. Self-administration took place in operant conditioning chambers (Med Associates, St. Albans, VT, USA) in sound-attenuating, ventilated cubicles. Each chamber was provided with two retractable levers, a liquid receptacle between them, two stimulus lights, and a sonalert above each lever. The receptacle was connected to an external syringe pump via polyethylene tubing. Presses on the right (active) lever activated the pump to deliver 0.1 ml of 10% alcohol and simultaneously turned on a 3-s light and tone. Presses on the left (inactive) lever were recorded but not reinforced.

*2-bottle choice and Operant Training.* Male Wistar rats first received 7 days of 24-h access to a 2-bottle choice [10% (v/v) alcohol vs. water] paradigm to promote alcohol taste familiarity and intake. Alcohol was then removed, and rats had the water bottle only for 2 days. Next, rats were trained to lever-respond for alcohol 10% (v/v) under an FR1 schedule in a 12-h operant session. Twenty-four hour later, self-administration continued in a 60-min session on Day 3, followed by 30-min sessions for the next two days. On Day 5, an inactive lever was added; responses were recorded but not reinforced. From this point onward, both levers were available during 30-min daily sessions.

*Fixed ratio (FR) 1 (FR1) schedule of reinforcement and chronic intermittent alcohol vapor exposure.* After 10 self-administration sessions, rats were assigned to air-exposed (non-dependent) or alcohol-vapor–exposed (dependent) groups based on active-lever responding and alcohol intake. Dependent rats underwent chronic intermittent alcohol vapor exposure (14 h ON/10 h OFF; target BAL 150–250 mg/dL) for 4 weeks. Alcohol self-administration then resumed in 30-min sessions three times weekly during acute withdrawal (6–8 h after vapor OFF), with non-dependent rats tested on the same schedule. After stabilization, CYM5442 (0, 2.5, 5, 10 mg/kg, IP) was administered 60 min before a 30-min FR1 test session in both groups.

*Progressive ratio (PR) schedule of reinforcement.* After a few sessions of regular self-administration, non-dependent and dependent rats were tested under the progressive ratio (PR) schedule of reinforcement. Under this conditions, the response requirement necessary to receive a drop of alcohol increased progressively according to Richardson and Roberts (3). The breakpoint (BP) value was defined as the last ratio attained by the rat over the 60 min session. CYM5442 was administered IP 60 min before the start of the PR test at the doses of 0, 5, and 10 mg/kg.

*Extinction and cue-induced Reinstatement of alcohol seeking behavior.* Rats underwent five consecutive 30-minute extinction sessions during which responses on the active lever were not reinforced, and the audiovisual cue previously paired with alcohol availability was turned off. The day following extinction, both non-dependent and dependent rats were administered either 0 or 10 mg/kg CYM5442 60 minutes before the Reinstatement test session, which lasted 60 minutes. Each rat was placed into the operant chamber and exposed to five non-contingent presentations of the alcohol-paired audiovisual cue immediately before the session began. After cues presentation, both levers were inserted into the chamber, and responses on the active lever resulted in the presentation of the cues without alcohol delivery.

*Spontaneous locomotor activity*

Acute effect of CYM5442 administration was assessed on spontaneous locomotor activity in male alcohol-naïve Wistar rats. Rats were randomly injected with either 0, 2.5, 5, and 10 mg/kg CYM5442 60 min before the start of the test and then placed into an arena provided with infrared photo beams (Med Associates, USA). The rat locomotor activity was measured over a 1 h session.

### 2. Total RNA Isolation and RNA-Sequencing.

CYM-treated CIE rats were sacrificed after the 30-minute alcohol self-administration session. Vehicle- treated naive controls (no self-administration) were sacrificed at the corresponding time. Microdissected tissue from the PFC was processed for total RNA isolation using the miRNeasy Kit (Qiagen, Redwood City, CA) and Zymo Purification Kit (Zymo Research, Irvine, CA, USA). Libraries were prepared with the VAHTS Universal V10 RNA-seq Library Prep Kit (stranded) for Illumina (Vazyme, San Diego, CA, USA). Poly(A)-based mRNA enrichment libraries were subsequently sequenced on NovaSeq6000 (Illumina) at 30M reads target coverage (150 bp paired-end reads).

*Gene expression measures and Gene Set Enrichment Pathway Analysis.*

All fastq files raw data passed the quality control by fastQC. *Fastp* software was employed to filter poor quality reads and remove Illumina adapters. Transcriptome mapping and annotation were carried out using Hisat2, Samtools and FeatureCounts with the current rat genome reference GRCr8 (GCF_036323735.1). Differential expression analysis was then performed using Deseq2. GSEA prerank was used for pathway analysis with ranking being -log10(pval)*sign(log2FC) from DEA.

### 3. CB1 and $S1P_1$ activity of CYM-5442.

CB1 activity of CYM-5442 and S1P1 receptor was tested by Multispan (Hayward, California) in MULTISCREEN™ CHO-K1 Cell Line Stably Expressing Human CB1 Receptor and MULTISCREEN™ CHO-K1 Cell Line Stably Expressing Human S1P1 Receptor using the MULTISCREEN™ cAMP 1.0 No Wash Assay Kit (Multispan, Inc.). $Ga_i$ Cyclic AMP (cAMP) Assay: Cells were thawed and plated in assay buffer at desired concentrations. cAMP assays were performed according to the manufacturer's protocol using MULTISCREEN™ TR-FRET cAMP 1.0 No Wash Assay Kit. In agonist mode, cells were treated with test compounds and control agonist ligands and incubated for 20 minutes at 37°C. In antagonist mode, cells were preincubated with test compounds for 15 minutes at room temperature prior to the addition of $EC_{80}$ concentration of control ligand, after which the cells were incubated for 20 minutes at 37°C. The reaction was terminated by sequentially adding MULTISCREEN™ 650-labeled cAMP and MULTISCREEN™ Eu-labeled anti-cAMP antibody in lysis buffer. The plate was then incubated at room temperature for 30 minutes before reading fluorescent emissions at 620 nm and 665 with excitation at 314 nm on FlexStation III (Molecular Devices).

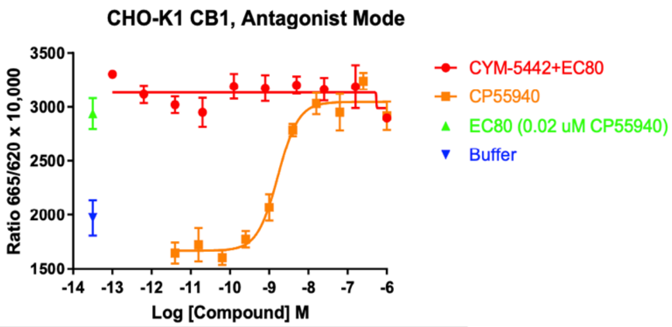


| | LogEC50 | HillSlope | EC50 |
|---|---|---|---|
| CP55940 | -8.778 | 1.651 | 1.666e-009 |

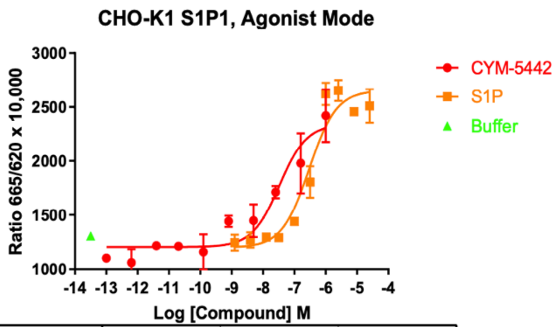


| | LogEC50 | HillSlope | EC50 |
|---|---|---|---|
| CYM-5442 | -7.478 | = 1 | 3.327e-008 |
| S1P | -6.528 | = 1 | 2.963e-007 |

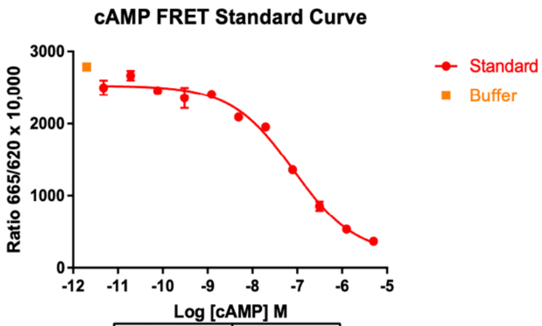


| | |
|---|---|
| LogEC50 | -7.088 |
| HillSlope | -0.6279 |
| EC50 | 8.173e-008 |

**Supplemental Fig. 1 Lack of CB1 activity of CYM5442.**
CYM5442 showed no against activity at the CB1 receptor, whereas CYM5442 displayed agonist activity for S1P1. Control agonists CP55940 and S1P showed dose-dependent inhibition of forskolin stimulated intracellular cAMP in the CB1 and S1P1 cells, respectively, with expected EC50 values.

### 4. References


1. Rhodes JS, Best K, Belknap JK, Finn DA, Crabbe JC. 2005. Evaluation of a simple model of ethanol drinking to intoxication in C57BL/6J mice. Physiol Behav 84:53-63.
2. Vendruscolo LF, Roberts AJ. 2014. Operant alcohol self-administration in dependent rats: focus on the vapor model. Alcohol 48:277-86.
3. Richardson NR, Roberts DC. 1996. Progressive ratio schedules in drug self-administration studies in rats: a method to evaluate reinforcing efficacy. J Neurosci Methods 66:1-11.
